# MITF regulates IDH1 and NNT and drives a transcriptional program protecting cutaneous melanoma from reactive oxygen species

**DOI:** 10.1101/2023.11.10.564582

**Authors:** Elisabeth Roider, Alexandra I.T. Lakatos, Alicia M. McConnell, Poguang Wang, Alina Mueller, Akinori Kawakami, Jennifer Tsoi, Botond L. Szabolcs, Anna A. Ascsillán, Yusuke Suita, Vivien Igras, Jennifer A. Lo, Jennifer J. Hsiao, Rebecca Lapides, Dorottya M.P. Pál, Anna S. Lengyel, Alexander Navarini, Arimichi Okazaki, Othon Iliopoulos, István Németh, Thomas G. Graeber, Leonard Zon, Roger W. Giese, Lajos V. Kemeny, David E. Fisher

## Abstract

Microphthalmia-associated transcription factor (MITF) plays pivotal roles in melanocyte development, function, and melanoma pathogenesis. MITF amplification occurs in melanoma and has been associated with resistance to targeted therapies. Here, we show that MITF regulates a global antioxidant program that increases survival of melanoma cell lines by protecting the cells from reactive oxygen species (ROS)-induced damage. In addition, this redox program is correlated with MITF expression in human melanoma cell lines and patient-derived melanoma samples. Using a zebrafish melanoma model, we show that MITF decreases ROS-mediated DNA damage *in vivo*. Some of the MITF target genes involved, such as *IDH1* and *NNT*, are regulated through direct MITF binding to canonical enhancer box (E-BOX) sequences proximal to their promoters. Utilizing functional experiments, we demonstrate the role of MITF and its target genes in reducing cytosolic and mitochondrial ROS. Collectively, our data identify MITF as a significant driver of the cellular antioxidant state.

**One Sentence Summary:** MITF promote melanoma survival via increasing ROS tolerance.

## INTRODUCTION

Melanoma is among the most common cancers in the northern hemisphere (*1*). Data from the US show an over 30-fold increase in melanoma incidence over the last century (*2*). Among US residents of European descent, the incidence of melanoma is about three times higher than in Asians and about 15 times higher than in individuals of South American or African origin (*3*). Because variations in melanin levels influence melanoma risk, the different effects of reddish-yellow pheomelanin and brown-black eumelanin on melanoma risk have been studied (*4*, *5*). Previously, we reported that pheomelanin synthesis promotes melanoma formation in a UV radiation-independent context along with significantly higher oxidative DNA and lipid peroxidation damage in *Mc1r* deficient, red-haired mice compared to genetically matched black (*Mc1r*-wildtype), albino, or combination red-albino mice (*5*). These findings were subsequently corroborated in humans as well (*6*) and highlight the importance of oxidative damage in the pathogenesis of melanoma, even independently of UV radiation.

Skin color is influenced by many genes; however, the microphthalmia-associated transcription factor (MITF) plays a master regulatory role in controlling skin pigmentation. MITF has several different isoforms with unique tissue-specific expression. The m-MITF isoform is uniquely expressed in the melanocyte lineage, and is subject to cAMP-mediated signal regulation of expression downstream of MC1R, the receptor whose loss-of-function is associated with red hair and pale skin (phototype 1). (*2*)

MITF belongs to a family of transcription factors with basic helix-loop-helix leucin zipper structures, which enable them to bind directly to canonical enhancer box (E-BOX) sequences (CA[T/C]GTG). Through binding to these sequences, M-MITF directly activates the transcription of hundreds of genes in the melanocytic lineage, and controls melanocyte proliferation, differentiation, survival, pigment production and a variety of additional processes. MITF induces pigmentation by acting as a transcription factor of TYR and numerous additional pigmentation genes including, but not limited to, TYRP1, DCT, PMEL, and MLANA (7–9). Other genes involved in melanocyte survival (e.g., BCL2, BCL2A1) and proliferation (e.g., CDK2) have also been identified as MITF target genes, amongst many other genes (10–12). Importantly, the production of eumelanin (brown pigment) is thought to shield the melanocytes and keratinocytes from UV damage by buffering the accumulation of UV-induced reactive oxygen species (ROS) during pigment production, as well as protecting cells from the UV-independent, pro-oxidant effects of pheomelanin (13). MITF has been shown to transcriptionally regulate PGC1α (PPARGC1A gene), a key protein that regulates cellular redox programs (14, 15). We hypothesized that, in addition to controlling the synthesis of the antioxidant eumelanin, MITF might promote ROS clearance by contributing to the regulation of cellular ROS homeostasis. Here, we show that MITF drives a global ROS clearance program in melanoma, by transcriptionally regulating multiple redox genes that contribute to the regulation of cellular ROS defense mechanisms.

## RESULTS

### MITF regulates genes involved in oxidative-reductive processes in melanoma

We first took an unbiased approach to investigate biological processes that MITF target genes are involved in. We performed gene ontology analysis (DAVID) (*16*, *17*) on genes that were significantly downregulated by at least 1.5 fold (log) in MALME-3M melanoma cells after MITF knockdown (*18*). We used REVIGO (*19*) to remove redundant gene ontology terms and visualize the significantly enriched gene sets. As expected, genes involved in regulation of cell survival and pigmentation were among the most highly enriched biological processes (Fig. 1a, Table S1). This is in line with previous observations regarding the roles of MITF in protecting the melanocytic lineage from apoptosis (*12*) and regulating enzymes necessary for melanin production. A gene set involved in the regulation of oxidation-reduction (redox) processes was also significantly enriched (Fig. 1A), suggesting that MITF might control a transcriptional program regulating cellular reactive oxygen production or elimination. Similarly, before removing redundant gene sets with REVIGO, gene ontology analysis by DAVID revealed that multiple gene sets related to regulation of redox processes, glutathione metabolism, and response to ROS (Fig. S1A, Table S2), are enriched in the downregulated genes after MITF knockdown. In an alternative pathway analysis using ToppCluster, we identified genes with NAD-binding and oxidoreductase activity that were significantly enriched among genes downregulated by MITF knockdown. These results collectively suggest that MITF might regulate a redox program in melanoma. Next, we performed another unbiased analysis using transcriptional profiles of 88 short-term melanoma cultures and investigated the functions of genes enriched in low vs. high MITF melanomas by using Gene Set Enrichment Analysis (GSEA) (*20*) with the Molecular Function gene set database. In this analysis, weighted statistic use ensures that poorly expressed genes and genes with low variance do not contribute to a positive enrichment score, which reflects the degree of overrepresentation of a gene set at the top or bottom of a ranked gene list. We observed that melanomas with high MITF were enriched in genes encoding proteins with oxidoreductase activity (Fig 1c, Fig. S1B, C). The observation that MITF expression across cell lines might be a predictor of differential redox activity further highlights the potential role of MITF in regulating cellular ROS.

**Fig. 1.**
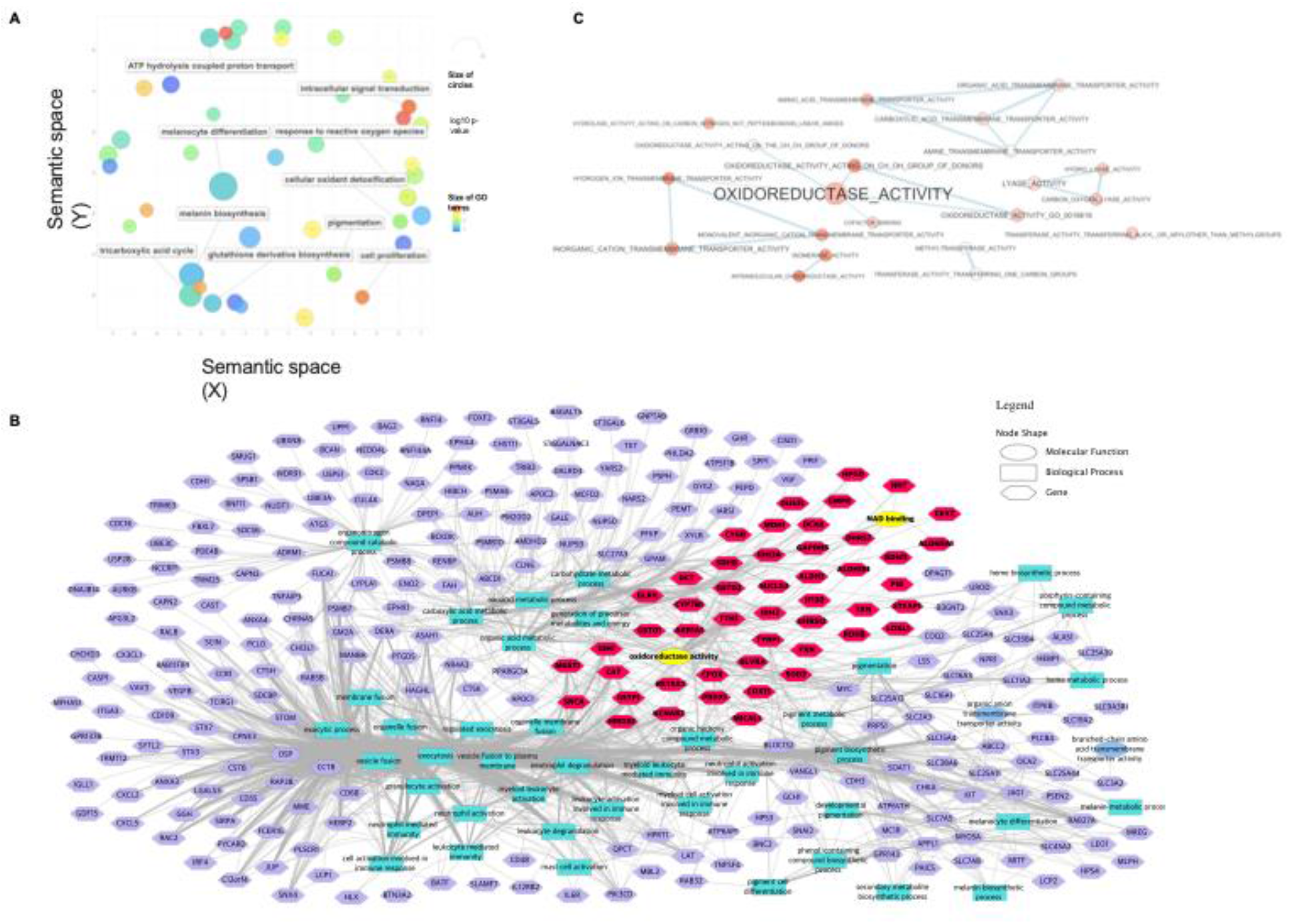
MITF^†^ regulates genes involved in oxidative-reductive processes in melanoma. **(A)** Gene ontology analysis (DAVID followed by REVIGO, for details see Methods) of downregulated genes after MITF knockdown in MALME-3M cells. **(B)** Pathway enrichment analysis of GO:Molecular functions and GO:Biological processes gene sets of downregulated genes after MITF knockdown in MALME-3M cells using graphic networks. NAD^‡^ binding and oxidoreductase activity are highlighted in yellow for clarity along with their associated genes that are highlighted in orange. (**C**) Oxidoreductase activity gene set is enriched in high MITF melanoma cell lines in short-term melanoma cultures. GSEA results were visualized by Cytoscape. † MITF: Microphthalmia-associated transcription factor ‡ Nicotinamide adenine dinucleotide

### MITF protects melanoma cells from oxidative stress *in vitro*

To functionally validate the bioinformatic findings, we next investigated the role of MITF in protecting melanoma cell lines from oxidative stress *in vitro*. Cytosolic and mitochondrial ROS levels were measured by DCFDA and Mitosox fluorescent dyes, respectively, using flow cytometry before/after knockdown of MITF in UACC257 and SKMEL5 melanoma cells (Fig. 2a). Confirmatory experiments were performed using confocal microscopy with UACC257 (Fig. 2b) and SKMEL5 (Fig. 2c) melanoma cells. As reduced glutathione (GSH) can be a key thiol antioxidant and a major detoxification agent in cells (*21*), we measured the effect of siMITF on GSH levels in UACC257 and SKMEL5 cells (Fig. 2d) and found that knocking down MITF decreased GSH levels significantly. To corroborate these findings, we analyzed the correlation of MITF with reduced glutathione levels in CCLE (*22*) (Fig. 2e) and in an independent previously published dataset where reduced glutathione levels were directly measured in human melanoma samples that were also transcriptionally profiled (*23*) (Fig. 2f). MITF significantly and positively correlated with reduced glutathione levels in both datasets, suggesting that it potentially plays a functional role in regulating cellular glutathione levels in melanoma.

**Fig. 2.**
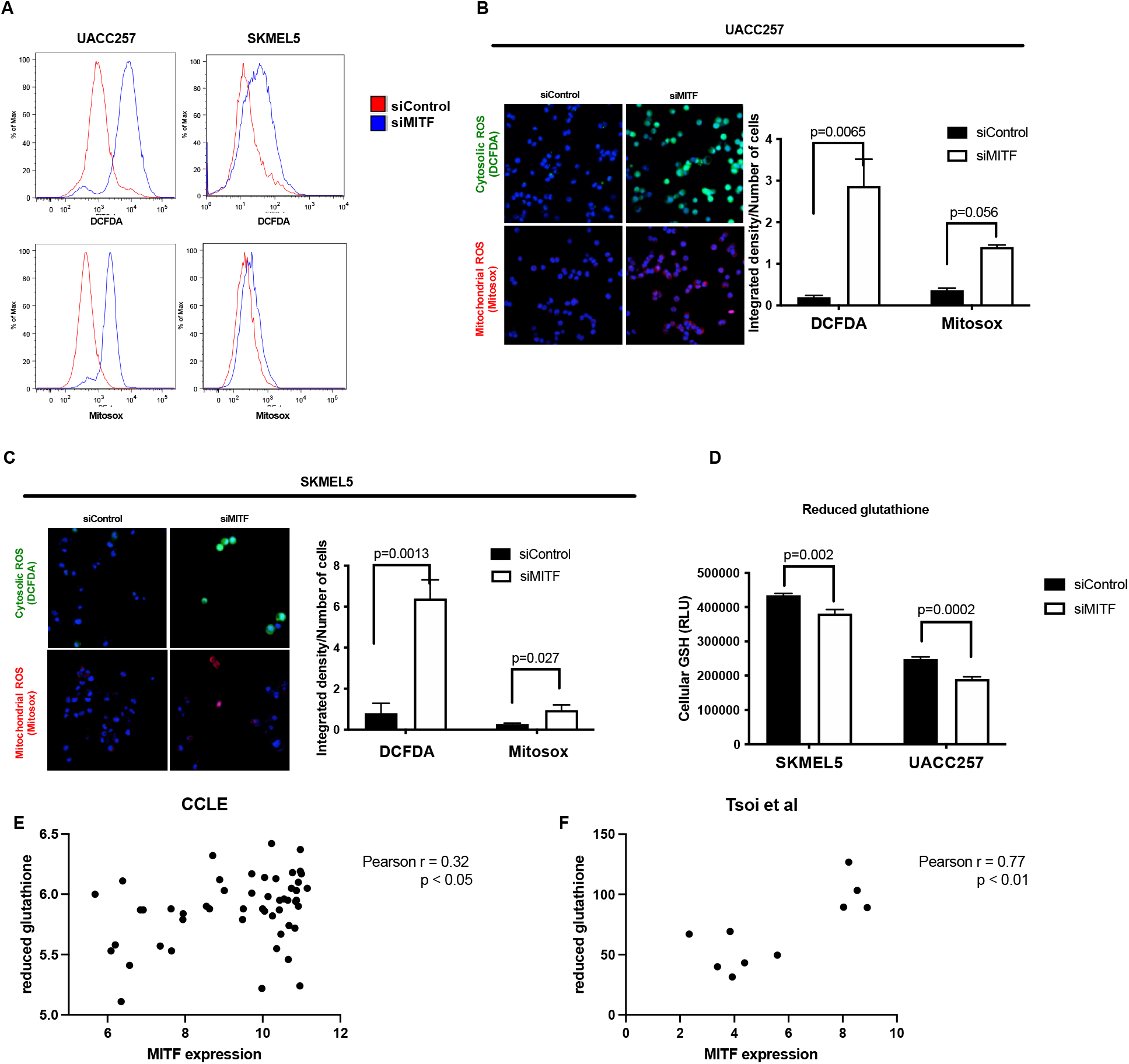
MITF^†^ protects melanoma cells from oxidative stress *in vitro*. (**A**) Representative flow plots for cytosolic ROS^‡^ levels measured by DCFDA fluorescent dye and mitochondrial ROS levels measured by Mitosox fluorescent dye in UACC257 melanoma cells after knockdown of MITF. (**B**) Confocal images of cytosolic ROS levels measured by DCFDA fluorescent dye and mitochondrial ROS levels measured by Mitosox fluorescent dye in UACC257 melanoma cells after knockdown of MITF. Representative images (left panel) and quantification of three independent experiments (right panel) are displayed. (**C**) Confocal images of cytosolic ROS levels measured by DCFDA fluorescent dye and mitochondrial ROS levels measured by Mitosox fluorescent dye in SKMEL30 melanoma cells after knockdown of MITF. Representative images (left panel) and quantification of three independent experiments (right panel) are displayed. (**D**) Reduced glutathione (GSH) was measured in UACC257 and SKMEL5 melanoma cells after siRNA silencing of *MITF*. Reduced glutathione levels significantly correlate with *MITF* expression in melanoma cell lines in CCLE (**E**) and in patient derived melanoma cells (**F**). † MITF: Microphthalmia-associated transcription factor ‡ Reactive oxygen species

### MITF directly induces expression of *IDH1* and *NNT*, which protect melanoma cells from oxidative stress

To further investigate the genes involved in the redox program regulated by MITF, we first identified potential MITF target genes involved in regulating redox processes. For this analysis, we examined the genes that were significantly downregulated by at least 1.5-fold after MITF knockdown in MALME-3M cells and selected genes that were involved in redox pathways (a complete list of the 47 genes selected and their corresponding gene ontology (GO) pathway gene sets are in Table S3). Next, we investigated the relationship between the expression of these genes and MITF expression across 61 melanoma cell lines in the CCLE database (Fig. 3a). We observed that most of the genes displayed a significant positive correlation with MITF. These results collectively suggest that MITF directly or indirectly regulates multiple genes involved in regulation of cellular ROS. To investigate the potential MITF target genes further, we used two publicly available MITF ChIP-seq datasets and analyzed regions in the proximity of each gene’s promoter sequences for MITF occupancy. Most of the genes had MITF occupancy surrounding their promoter regions, suggesting that MITF may directly regulate the transcription of these genes (Fig. 3a, genes in bold type). We also found potential direct binding sites for MITF surrounding these genes by identifying consensus binding E-BOX sequence elements for MITF, in close proximity to their promoter regions. However, MITF occupancy was not detected in close proximity to the promoter regions of one third of the genes, suggesting that MITF may indirectly regulate transcription of these genes by either binding to distal enhancer elements or by modulating the expression of other transcription factors. Collectively, these results suggest that the MITF-driven redox program is due to both indirect and direct transcriptional regulation by MITF.

**Fig. 3.**
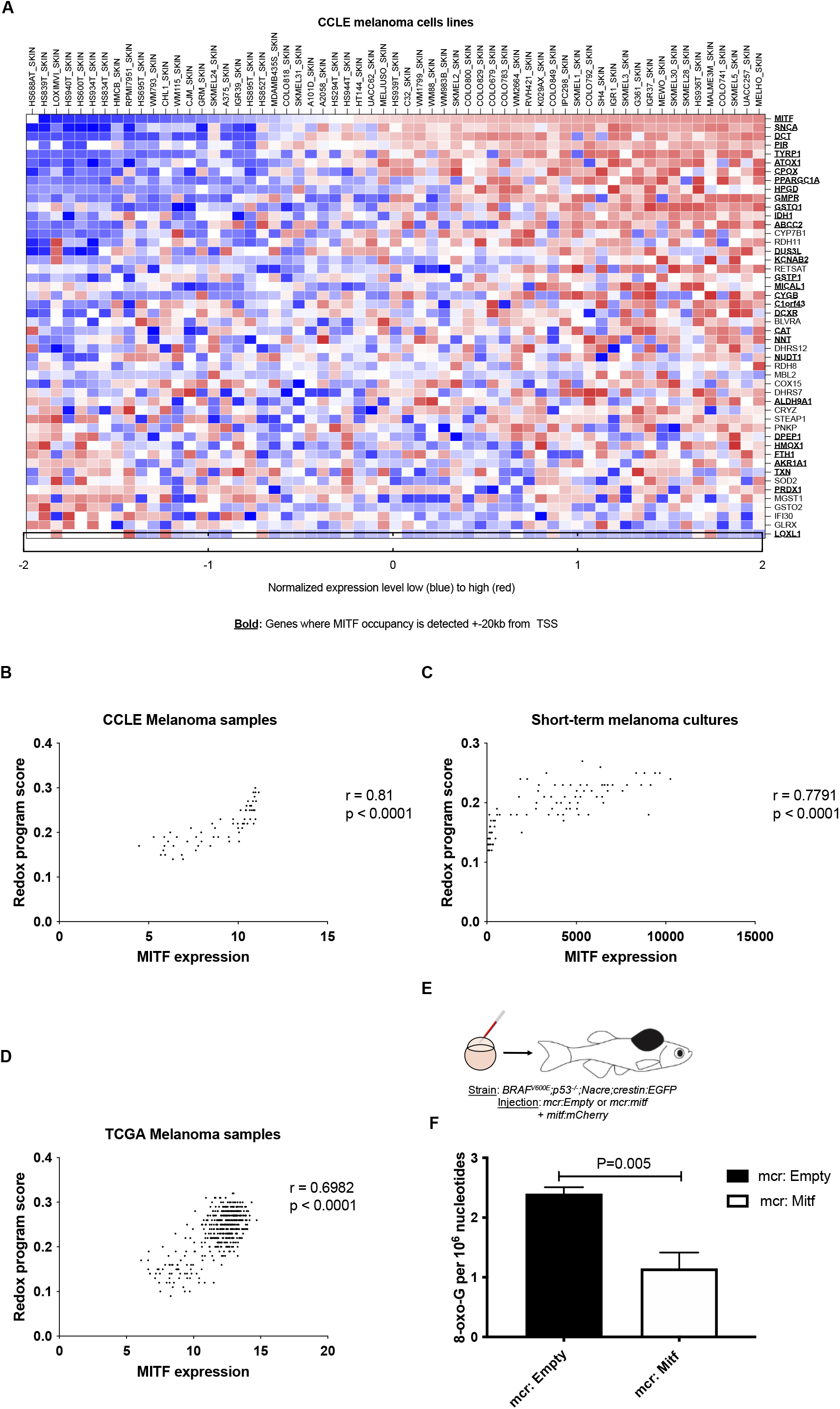
MITF^†^ directly regulates a redox program in melanoma in vivo. (**A**) Correlation of downregulated genes after *MITF* knockdown that are annotated with regulation of cellular redox levels across 61 melanoma cell lines in the CCLE microarray dataset. MITF occupies promoter or enhancer regions of most of these genes (defined as MITF ChIP peaks within ± 20kb of a transcription start site (TSS), indicated with bold) based on a publicly available MITF ChIP-sequencing dataset and a ChIP-ChIP dataset. The genes displayed in panel **a** were used to define an MITF-driven redox program, which was scored using the singscore algorithm in human patient tumor samples from TCGA. Singscore algorithm defined MITF-driven redox program significantly correlates with MITF mRNA levels across melanoma cell lines (CCLE, B), short-term melanoma cultures (Lin dataset, C) and. 474 human melanoma samples (TCGA, D). (**E**) *mitfa:BRAF^V600E^*;*p53^−/−^*;*mitfa^−/−^;crestin:EGFP* zebrafish were injected at the single cell stage with a MiniCoopR plasmid that rescues melanocytes and expresses human MITF under control of the zebrafish *mitfa* promoter (*mcr:MITF*) or with the same plasmid without the human MITF expression cassette (*mcr:Empty*), along with a plasmid expressing the mCherry marker from the zebrafish *mitfa* promoter (*mitf:mCherry*). HPLC-based redox measurements in melanoma tumors from zebrafish re-expressing human MITF (*mcr:MITF*) or control plasmid (*mcr:Empty*) at 2 weeks post-fertilization show decreased 8-oxoG levels in MITF overexpressing zebrafish tumors (**F**). † MITF: Microphthalmia-associated transcription factor

### MITF protects melanoma from oxidative damage *in vivo*

Our results suggest that MITF directs a transcriptional program that protects melanoma cells from oxidative stress *in vitro*. Next, we aimed to investigate the function of this program *in vivo*. First, we defined an MITF-driven redox program based on the genes annotated to redox processes whose expression in MALME-3M cells changes significantly after *MITF* knockdown (as discussed above and displayed in Fig. 3a and Table S3). We used a rank-based, single-sample gene set scoring method (singscore algorithm) (*24*) to assign an MITF-driven redox program score for melanoma cell lines (from CCLE), short-term melanoma cultures (*25*, *26*), and patient samples from TCGA (Cancer Genome Atlas, 2015). As expected, MITF expression correlated significantly with its redox program score among both in vitro datasets (Fig. 3b, c). Importantly, we also observed a significant correlation across 474 patient melanoma samples from TCGA (Fig. 3d), suggesting that the MITF-driven redox program is relevant in patient biopsies despite the presence of non-malignant stromal and immune cell populations.

In order to investigate whether MITF controls ROS levels *in vivo,* we examined a zebrafish melanoma model. We used the MiniCoopR system to overexpress human MITF in melanoma-prone zebrafish (Fig 3e). *mitfa:BRAF^V600E^*;*p53^−/−^*;*mitfa^−/−^*;*crestin:EGFP* zebrafish lack melanocytes due to a deletion in the zebrafish *mitfa* gene. These fish were injected at the single cell stage with a MiniCoopR plasmid, which contains both an *mitfa* minigene to rescue melanocyte development, and an *mitfa:MITF* cassette which expresses the human *MITF* gene under control of the zebrafish *mitfa* promoter (*mcr:MITF*) (*27*). This allowed us to ensure that every rescued melanocyte present in the zebrafish is overexpressing MITF. These fish were compared to zebrafish injected with a MiniCoopR plasmid expressing only the *mitfa* minigene (*mcr:Empty*), which results in melanocyte rescue with no MITF overexpression. A separate plasmid with mCherry expression driven by the *mitfa* promoter was co-injected to visualize the rescued melanocytes. We used *crestin:EGFP* as a marker of the embryonic neural crest, which is specifically reactivated in melanomas (*28*) (Fig. S2A). We observed that MITF overexpression in zebrafish resulted in a significant increase in the induction of *crestin:EGFP* at six weeks post-fertilization (Fig. S2B) and had significantly accelerated tumor onset compared to *mcr:Empty* controls (Fig. S2C), as expected due to the oncogenic properties of MITF. To investigate whether MITF overexpression influences ROS-mediated DNA damage *in vivo*, we assessed levels of the DNA oxidation product 8-oxoG. 8-oxoG levels (Fig. S3) were significantly lower in zebrafish melanoma tumors overexpressing MITF compared with control tumors (Fig. 3f). These data indicate that high MITF levels promote melanoma formation and, consistent with the in vitro studies above, decrease ROS levels *in vivo*.

### MITF directly regulates transcription of isocitrate dehydrogenase 1 (IDH1) and nicotinamide nucleotide transhydrogenase (NNT)

To further validate direct transcriptional regulation of some of the candidate genes by MITF, first we investigated the mechanism of regulation of *NNT* and *IDH1* by MITF, as both have E-BOX sequences in the proximity of their transcription start sites, and are involved in nicotinamide adenine dinucleotide phosphate (NADPH) and GSH metabolism. The enzyme *NNT*, which catalyzes the transfer of reducing equivalents from NADH to NADPH, is a known regulator of mitochondrial redox levels located in the inner mitochondrial membrane. NNT is regarded as a major source of NADPH and reduced glutathione in mitochondria (*29*). IDH1 is an enzyme with similar function with regard to NADPH formation, but is located in the cytosol (*30*). Given the presence of E-BOX sequences and MITF occupancy in the proximity of promoter regions and their correlation with MITF expression in the CCLE microarray (Fig. 3a) and CCLE RNA-seq datasets (Fig. 4a, b), we hypothesized that MITF directly regulates the transcription of *NNT* and *IDH1*.

**Figure 4.**
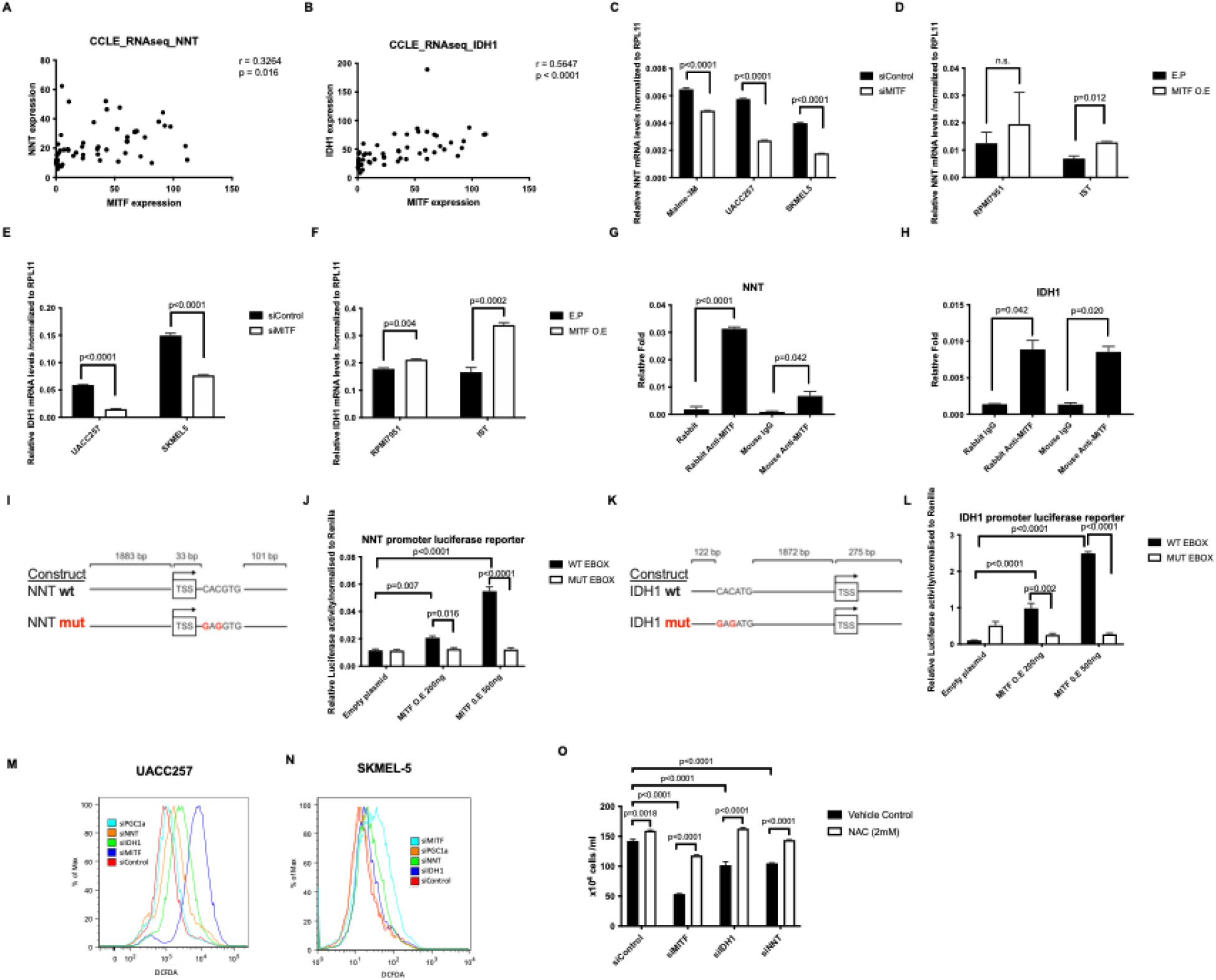
MITF^†^ directly targets IDH1 and NNT and they protect melanoma from oxidative damage Similar to the microarray dataset, we found significant correlations of MITF with NNT^‡^. (**A**) and *IDH1*^§^ (**B**) across 54 melanoma cell lines in the CCLE RNA-sequencing dataset. *NNT* mRNA levels following knockdown (**C**) or overexpression (**D**) of *MITF* (48 h) in human melanoma cells. IDH1 mRNA levels following knockdown (**E**) or overexpression (**F**) of *MITF* (48h) in human melanoma cells. ChIP with polyclonal and monoclonal anti-MITF antibodies in UACC257 melanoma cells. Precipitated DNA was amplified using primers surrounding an MITF binding site within the *NNT* (**G**) or *IDH1* (**H**) promoter region and compared to rabbit/mouse IgG control. Effects of *MITF* overexpression on luciferase reporter activity driven by the NNT (**I, J**) *IDH1* promoter (**K, L**) with wild type (WT) or impaired (MUT) E-box sequences in UACC257 cells. Representative flow plots for cytosolic ROS levels measured by DCFDA fluorescent dye after knockdown of *MITF*, *IDH1*, *NNT*, or *PGC1α* in UACC257 melanoma cells (**M**) and in SKMEL30 melanoma cells(**N**). *MITF*, *IDH1*, or *NNT* silencing induced ROS dependent melanoma death that is rescuable by 2 mM NAC (N-Acetyl-L-Cystein) in UACC257 melanoma cells (**O**). Values represent SD of three independent experiments performed. P values in *m* and *n* indicate significance compared to siControl. † MITF: Microphthalmia-associated transcription factor ‡ Nicotinamide nucleotide transhydrogenase § Isocitrate dehydrogenase 1

*In vitro* experiments confirmed the direct effect of MITF on *NNT* and *IDH1* mRNA levels. First, mRNA levels of *NNT* were downregulated and upregulated after knockdown (Fig. 4c) and overexpression (Fig. 4d) of MITF in UACC257 melanoma cells, respectively. Similar effects were found for the mRNA levels of *IDH*1 following knockdown (Fig. 4e) and overexpression (Fig. 4f) of MITF in UACC257 cells. In order to demonstrate a direct transcriptional role of MITF on *IDH1* and *NNT*, chromatin immunoprecipitation (ChIP) with anti-MITF antibodies in UACC257 melanoma cells was performed. Using ChIP-qPCR, we observed that MITF occupies *NNT* (Fig. 4g) or *IDH1* (Fig. 4h) promoter regions. Next we used luciferase reporters to express the promoter regions of *NNT* and *IDH1*, both of which contain the canonical E-box sequence in close proximity to their transcription start sites (illustrated in Fig. 4i,k). MITF overexpression dose-dependently promoted luciferase activity of both *NNT* and *IDH1* promoters (Fig. 4j,l), but not when the E-BOX sequence was mutated to prevent MITF binding. These results suggest that MITF binding to the *IDH1* and *NNT* promoters directly promotes their transcription.

Next, we aimed to functionally investigate the roles of NNT and IDH1 in regulating cellular ROS in melanoma. In addition, the effect of PGC1α, a known regulator of oxidative stress and reduced glutathione, cystathionine, and 5-adenosylhomocysteine levels (*31*) was investigated. Knockdown of *MITF*, *IDH1*, or *NNT* in UACC257 cells (Fig. 4m) and *MITF* or *IDH1* in SKMEL5 cells (Fig. 4n) resulted in significantly increased cytosolic oxidative stress levels, as shown by flow cytometry (left panels) measuring cytosolic ROS levels by DCFDA fluorescent dye. To examine how this increase in ROS affects survival of melanoma cells, we measured viability of UACC257 melanoma cells upon *MITF*, *NNT*, or *IDH1* knockdown, in the presence or absence of the thiol antioxidant N-Acetyl-L-Cysteine (NAC). We observed that knockdown of *MITF*, *NNT*, or *IDH1* decreased cell viability significantly, and all could be significantly rescued by adding NAC (Fig. 4o). Of note, *MITF* suppression had the greatest impact on ROS induction (and viability) with *NNT* or *IDH* showing intermediate effects—consistent with the NNT/IDH roles as contributors to MITF’s overall effects, while further suggesting that additional MITF-regulated genes also likely contribute to both the redox and viability phenotypes.

## DISCUSSION

Although elevated ROS levels can contribute to tumorigenesis and early tumor growth, rapidly proliferating cancer cells must attenuate excessive ROS levels to survive (*5*, *32*). NRF2 is widely recognized as a major transcriptional regulator of the anti-oxidant state in cells. Induction of eumelanin synthesis has been shown to exert anti-oxidant effects (*13*), however, we provide evidence here that MITF is capable of playing a lineage-specific role in contributing to reduction of oxidative stress through transcriptional activation of multiple genes. Recent evidence has indicated that MITF can also induce expression of proteins with antioxidant properties, such as the mitochondrial biogenesis regulator PGC1α (*31*). This evidence prompted us to apply an unbiased approach to elucidate the extent of MITF-regulated expression of redox-related genes in melanoma cells. Genome-wide analysis of gene expression in melanoma cells, with or without *MITF* knockdown, and analysis of MITF binding regions confirmed the known impact of MITF on genes involved in pigmentation (*18*), DNA replication, mitosis, and DNA repair (*33*), and also revealed robust enrichment of genes involved in responses to oxidative stress. Forty-six independent redox-related genes including *PGC1α* (*PPARGC1A*) were identified as direct or indirect targets of MITF; and most of them had not previously been identified as target genes of MITF.

Expression levels of most of the redox-related genes were highly correlated with MITF across multiple short-term melanoma cultures, melanoma cell lines, and patient tumor samples, accentuating the robustness of the MITF-driven redox program in melanoma. *In vitro* data from *MITF* knockdown in different human melanoma cell lines confirmed the role of MITF in limiting mitochondrial ROS (*15*, *34*) and revealed more robust effects of MITF on cytosolic ROS. Moreover, MITF was found to decrease levels of 8-oxo-G levels, a marker of DNA oxidation, oxidation *in vivo* when overexpressed in zebrafish melanoma tumors. These findings are in line with our previous observation showing that red-hair, pheomelanin-rich mice display increased ROS-mediated DNA damage, potentially contributing to the carcinogenic risk red-haired individuals carry (*5*).

Nicotinamide adenine dinucleotide (NAD+/NADH) and nicotinamide adenine dinucleotide phosphate (NADP+/NADPH) are major determinants of the cellular redox state (*35*). In order to understand how MITF may regulate the cellular redox system, the roles of two selected MITF-dependent NADPH replenishers, NNT and IDH1, were investigated. To confirm the direct regulatory impact of MITF on NNT and IDH1, chromatin immunoprecipitation and luciferase reporter experiments were performed. Whereas the interplay between MITF and individual redox genes such as *HIF1α* (*14*), *PGC1α* (*15*) and *APEX1* (*36*) has been shown before, our study revealed the extent of how deeply MITF is involved in the redox metabolism of human melanoma.

Melanomas are characterized by high MITF-expressing and low MITF-expressing tumors (*37*), and subsets of MITF-high and MITF-low cells appear to exist in virtually all melanoma tumors based upon single cell analyses (*38*). A fluid state between MITF-high and MITF-low melanomas, presumably at least partially in response to different environmental triggers, has been proposed as a rheostat-like modulation model (*37*). MITF-high melanomas are more proliferative, differentiated, and less invasive than MITF-low melanomas. This is mainly due to the induction of various cell cycle and differentiation genes by MITF and consistent with its antioxidant properties, which are necessary to mitigate the increased ROS generated during energy production for rapid proliferation (*39*). Indeed, the antioxidant MITF target, PGC1α, has been shown to enhance proliferation of a subset melanoma cell lines in which it is expressed, while inhibiting metastasis of those cells (*31*, *40*).

The dedifferentiated state of melanoma has previously been associated with reduced glutathione levels and high sensitivity to the iron-dependent, lipid ROS-mediated ferroptotic cell death (*23*) . It is possible that MITF might directly regulate ferroptosis or other changes in cell metabolism in melanoma; however future studies are required to assess the role of MITF in regulating sensitivity to ferroptotic cell death.

In summary, beyond the essential role of MITF in melanoma survival and oncogenesis, we identified an MITF-regulated redox program with multiple new direct and indirect transcriptional targets that eliminate cellular ROS. Understanding of the basis of melanoma biology, and especially the differences between high and low MITF melanomas, may not only help in the design of tailored prevention strategies, but also lay the groundwork for future therapeutic directions.

## MATERIALS AND METHODS

### Cell culture

Human UACC257 and SKMEL5 melanoma cell lines were obtained from NCI and grown in DMEM or RPMI medium supplemented with 10% fetal bovine serum and 1% penicillin/streptomycin/L-glutamine.

### siRNA delivery

A single pulse of 10 nmol/L of siRNA was delivered to a 60% confluent culture by lipidoid transfection as described before (*12*). Lipidoid material was synthesized by reaction of 1,2-epoxydodecane with 2,2’-diamino-Nmethyldiethylamine in a glass scintillation vial for 3 days at 90°C. Following synthesis, the reaction mixture was characterized by MALDI-TOF mass spectroscopy to confirm mass of expected products. Reaction product was used for transfection without further purification. Lipidoid was dissolved in 25 mM NaOAc buffer (pH ∼5.2) and added to a solution of siRNA for complexation. Complexes of siRNA (final concentration of 25 nM) were plated in 96 well plates, followed by plating cells in growth medium as above. After 48-72 hours of transfection, total RNA or protein was harvested. The following siRNA pools were purchased from Dharmacon: siGENOME Human MITF SMARTpool (M-008674), siGENOME Human IDH1 SMARTpool (M-008294), siGENOME Human NNT SMARTpool (M-009809), siGENOME Human PPRGC1A SMARTpool (M-008294) and ON-TARGETplus non-targeting control pool (D-001810).

### RNA purification and quantitative RT-PCR

RNA was harvested from melanoma cells 48 hours after siRNA or overexpression vector transfection by using RNeasy Plus mini kit (Qiagen) according to the manufacturer’s instructions. RNA was harvested from mouse ear skin using TissueLyser II (Qiagen) and TRIzol (Life Technologies) according to the manufacturers’ instructions, followed by second purification using an RNeasy Plus mini kit (Qiagen). mRNA expression of melanocytic markers and PD-L1 was determined using intron-spanning primers with SYBR FAST qPCR master mix (Kapa Biosystems). qRT-PCR was performed with the following primers. Human RPL11: forward, GTTGGGGAGAGTGGAGACAG; reverse, TGCCAAAGGATCTGACAGTG. Human M isoform MITF: forward, CATTGTTATGCTGGAAATGCTAGAA; reverse, GGCTTGCTGTATGTGGTACTTGG; Human NNT: forward, AGCTCAATACCCCATTGCTG; reverse, CACATTAAGCTGACCAGGCA. Human IDH1: forward, GTCGTCATGCTTATGGGGAT; reverse, CTTTTGGGTTCCGTCACTTG. Human PPRGCA1: forward, CTGCTAGCAAGTTTGCCTCA; reverse, AGTGGTGCAGTGACCAATCA. Expression values were calculated using the comparative threshold cycle method and normalized to human RPL11 or mouse 18S RNA.

### Flow cytometry

MitoSOX (M36008) was obtained from Thermo Fisher Scientific and 2’,7’–dichlorofluorescein diacetate (DCFDA or H2DCFDA) was obtained from Abcam (ab113851) and used according to the manufacturers’ recommended protocols. Mean fluorescence was determined by FlowJo software (BD Biosciences) and normalized to vehicle-treated cells.

### Confocal microscopy and quantification

Adherent cells were cultured on a glass bottom dish and incubated according to the manufacturer’s protocols. The following settings were used: 5 μM MitoSOX Red (Thermo Fisher Scientific, M36008) in PBS/5% FBS at 37°C for 10 min or 2 μM H2-DCFDA (Thermo Fisher Scientific, C6827) in PBS/5% FBS at 37°C for 30 min, followed by washing with HBSS. Stained cells were analyzed by immunofluorescence imaging and normalized to cell numbers, which were detected by nuclear staining with 1 drop per mL Nucblue (Thermo Fisher Scientific, R37605) at 37°C for 15 min.

### Glutathione measurements

Cell lysates from equal numbers of cells were analyzed for glutathione using GSH/GSSG-Glo assays (Promega, V6611) according to the manufacturer’s protocol.

### Cell viability measurements and NAC rescue

Cell numbers were counted manually using trypan blue (Abcam, ab233465) and indicated cell lines were enriched with 2 mM N-Acetylcysteine (Sigma-Aldrich, 616-91-1).

### Lentivirus production and infection

Lentiviruses were produced as previously described (*41*). Cells were split 1 day before infection. Cells were centrifuged at 1,000 × g for 30 min in a suitable medium for each cell type with lentiviruses and polybrene (final concentration 8 μg/mL). On the second day of infection, the medium was replaced with fresh medium containing puromycin at a suitable concentration for each cell type. Cells were harvested at the indicated time points.

### Chromatin Immunoprecipitation (ChIP)

Human primary melanocytes were fixed with formaldehyde in PBS (1% final concentration) for 15 min at room temperature. Fixed cells were scraped with ice cold PBS containing protease inhibitor (Roche). 5 million cells were suspended in 500 μl SDS lysis buffer (50 mM Tris-HCl, pH 8.0, 10 mM EDTA, 1% SDS, protease inhibitor (Roche)). 50 μg of DNA/protein complexes were rotated for 10 min at 4°C, and sonicated by Bioruptor® (Diagenode) to yield DNA fragments around 500 base pairs. Samples were centrifuged to remove debris and supernatants were diluted 10 times with IP dilution buffer (0.01% SDS, 1.1% Triton X-100, 1.2 mM EDTA, 16.7 mM Tris-HCl, pH 8.0, 167 mM NaCl, protease inhibitor). To reduce background, samples were pre-cleared with 5 μg of normal rabbit IgG (Santa Cruz Biotechnology) and 80 μl of 50% protein A/G slurry containing 0.25 mg/mL sonicated salmon sperm DNA and 1 mg/mL BSA for 2 hours at 4°C. Antibodies were added to pre-cleared chromatin solution and incubated overnight at 4°C. Protein A/G slurry containing 0.25 mg/mL sonicated salmon sperm DNA and 1 mg/mL BSA were added to samples and incubated for 2 hours at 4°C. Immunocomplexes were washed twice with low salt buffer (0.1% SDS, 1% Triton X-100, 2 mM EDTA, 20 mM Tris-HCl, pH 8.1, 150 mM NaCl, protease inhibitor), twice with high salt buffer (0.1% SDS, 1% Triton X-100, 2 mM EDTA, 20 mM Tris-HCl, pH 8.1, 500 mM NaCl, protease inhibitor), once with LiCl buffer (0.25 M LiCl, 1% NP40, 1% sodium deoxycholate, 1 mM EDTA, 10 mM Tris-HCl, pH 8.1, protease inhibitor), and twice with TE containing protease inhibitor. Immunocomplexes were eluted from beads with 50 μl elution buffer (1% SDS, 10 mM DTT, 0.1 M NaHCO_3_, protease inhibitor) and rotated for 15 min twice at room temperature. Crosslinks were reversed overnight at 65°C. Proteins were digested by 1 hour incubation with proteinase K at 56°C. DNA was purified using a QIAquick PCR Purification Kit (Qiagen).

### Reporter assay

*IDH1* and *NNT* promoter sequences were amplified from genomic DNA of human primary melanocytes by PCR and cloned into pGL4.12 (Promega). Mutagenesis of E-boxes was performed using a QuikChange Site-Directed Mutagenesis Kit (Stratagene). Melanoma cells were transfected with combinations of reporter constructs: pRL-CMV (Promega) and either pCDH-Cuo-IRES-RFP or pCDH-Cuo-hMITF-M-IRES-RFP, with polyethylenimine. Two days after transfection, melanoma cells were harvested in Passive Lysis Buffer (Promega). Firefly and Renilla luciferase activities were measured by Dual-Luciferase Reporter Assay System (Promega) using FLUOstar Omega (BMG Labtech). Primer sequences for mutagenesis of the IDH1 promotor: forward, GGGAGAAGGTCAGCAGGAAACATCTCAGCAAAGGAATC; reverse, GATTCCTTTGCTGAGATGTTTCCTGCTGACCTTCTCCC. Primer sequences for mutagenesis of the *NNT* promotor: forward, CTAGCTAGCAGTCAGGGAGGGAGGAAAGAGTAGAA; reverse, GAAGATCTTTGGGCTGTGCCCTGAG.

### Identification of an MITF-related redox gene set

To identify this cluster of MITF-regulated redox genes, the oxidation-reduction process gene ontology gene set GO:0055114 was intersected with genes that are down-regulated at least 1.5-fold upon MITF knockdown in MALME-3M melanoma cells (*35*). Gene expression levels were obtained from 61 melanoma cells lines in the cancer cell line encyclopedia (CCLE) microarray dataset (*38*). Further analysis included testing of MITF ChIP peaks surrounding promoter or enhancer regions of these redox genes. MITF occupies promoter regions of some of these genes, based on a publicly available MITF ChIP-sequencing dataset (*33*) and a ChIP-ChIP dataset (*42*) via checking the presence of E-BOX sequences (CACGTG and CATGTG) that are consensus binding sites for MITF family-related transcription factors.

DAVID (*16*, *17*) was used to identify gene sets downregulated by MITF knockdown by at least 1.5-fold in the MALME-3M cell line dataset. REVIGO (*18*, *19*) with SimRel semantic similarity measure and with allowed similarity of 0,5 was used to remove redundant gene sets using the significantly enriched gene sets (p = 0.05) by DAVID. GSEA analyses were conducted using GSEA v2.07 (*20*, *43*)with default parameters. The singscore R/Bioconductor package was used to score the MITF-driven redox program in individual melanoma samples (*24*).

### Pathway analyses

For the alternative pathway analysis (Fig 1b), first, genes that were downregulated in MALME cells with at least 1.5-fold were selected and used as an input and were used with standard parameters, except for Bonferroni p-value cutoff being 0.2. The ToppCluster gene list feature analysis was used with default settings as an input to Cytoscape 3.6.1. Gene sets enriched in MITF high melanoma cell lines were identified by GSEA focusing on GO Molecular Function gene set database. Then EnrichmentMap in Cytoscape was used to visualize the results of gene set enrichment. The analysis was made with Cytoscape 3.9.1.

### Detection of 8-Oxoguanine (8-oxoG) by mass spectrometry

This was done as described elsewhere (*44*). Briefly, zebrafish melanoma DNA was purified further (including desalting) by spinning in an Amicon Ultra 0.5 mL Centrifuge Filter (regenerated cellulose 3000 NMWL); discarding the filtrate; adding 320 µL of water:acetonitrile, 9:1 (v/v); spinning and discarding; repeating four more times; rinsing the inner surface of the filter with 50 µL of water; reversing the filter; and centrifuging again to obtain 100 µL of water containing desalted DNA. The DNA solution was combined in a ratio of 1:1 (v/v) with 4-hydroxy-alpha-cyanocinnamic acid in 50% acetonitrile; MALDI-MS then was used to detect the nucleobase, and relative quantitation was achieved by comparing this peak height to the average heights for adenine and guanine.

### Expression of MITF in a zebrafish melanoma model

The human MITF gene was cloned into the MiniCoopR overexpression plasmid under the control of the *mitfa* promoter (*mcr:mitf*). This allows for the rescue of melanocytes in *nacre* (*mitfa^−/−^*) mutant zebrafish by overexpression of MITF in melanocytes (*27*). The *mcr:mitf* or *mcr:Empty* control plasmid together with *mitf:mCherry* plasmid were injected into *BRAF^V600E^*;*p53^−/−^*;*nacre*;*crestin:EGFP* zebrafish embryos at the single cell stage and incorporated into the genome using Tol2 transgenesis. Fish were imaged under a Nikon SMZ18 Stereomicroscope at 6 weeks of age and observed until 20 weeks of age for melanoma tumor formation. The zebrafish experiments performed in this study were in strict accordance with the recommendations in the Guide for the Care and Use of Laboratory Animals of the National Institutes of Health. The animal research protocol was approved by the Institutional Animal Care and Use Committee of Boston Children’s Hospital. All zebrafish used in this study were maintained and euthanized under the guidelines of the Institutional Animal Care and Use Committee of Boston Children’s Hospital.

### Statistical analysis

Statistical analyses were performed using GraphPad Prism 8. Single comparisons of two groups were analyzed by two-tailed Student’s t tests, correcting for multiple pairwise comparisons when applicable using the Holm-Šidák post-test. Comparisons of more than two groups with single independent and dependent variables were analyzed by one-way ANOVA with the Brown-Forsythe and Welch modification to account for different standard deviations and Dunnett’s correction for multiple pairwise comparisons. Multiple pairwise comparisons of two factor experiments across factors (Fig. 3J, K) were analyzed by two-way ANOVA with the Holm-Šidák correction for multiple pairwise comparisons. P values less than 0.05 were considered statistically significant.

## General

This work was conducted with support from Harvard Catalyst | The Harvard Clinical and Translational Science Center (National Center for Advancing Translational Sciences, National Institutes of Health Award UL 1TR002541) and financial contributions from Harvard University and its affiliated academic healthcare centers. The content is solely the responsibility of the authors and does not necessarily represent the official views of Harvard Catalyst, Harvard University and its affiliated academic healthcare centers, or the National Institutes of Health.

## Funding

ER acknowledges support from the Mildred Scheel Grant of the German Cancer Society and the Filling the Gap grant of the University of Zurich, Switzerland. L.V.K. is a recipient of the János Bolyai Research Scholarship of the Hungarian Academy of Sciences and the work was supported by the Hungarian National Research, Development and Innovation Office (OTKA FK138696), KIM NKFIA TKP-2021-EGA-05 and KIM NKFIA 2022-2.1.1-NL-2022-00005 grants. T.G.G. is supported by NIH P01 CA244118 and Melanoma Research Alliance 691165. DEF acknowledges support to his laboratory from NIH grants P01 CA163222, R01 AR072304, and R01 AR043369, as well as funding from the Dr. Miriam and Sheldon G. Adelson Medical Research Foundation and the Melanoma Research Alliance.

## Author contributions

Study concept (E.R., L.V.K., D.E.F.), methodology (E.R. A.I.T.L., L.V.K., D.E.F.), investigation (E.R., A.I.T.L., A.M.M., P.W., A.M., A.K., J.T., B.L.Sz., A.A.A., Y.S., V.I., J.A.L., J.J.H., R.L., D.M.P.P., A.S.L.), supervision (A.N., A.O., O.I., I.N., T.G.G., L.Z., R.W.G., L.V.K., D.E.F.), original draft writing (E.R., L.V.K., D.E.F.) and review and editing of the manuscript (A.I.T.L., R.L., A.S.L.).

## Competing interests

Dr. Fisher has a financial interest in Soltego, Inc., a company developing SIK inhibitors for topical skin darkening treatments that might be used for a broad set of human applications. Dr. Fisher’s interests were reviewed and are managed by Massachusetts General Hospital and Partners HealthCare in accordance with their conflict-of-interest policies. Dr. Roider is a shareholder and founder of Maximon AG and its holding ventures. All other authors declare no conflicts of interest. All other authors declare they have no competing interests.

## Data and materials availability

All data are available in the main text or the supplementary materials.

## Supplementary Materials

**Fig. S1.**
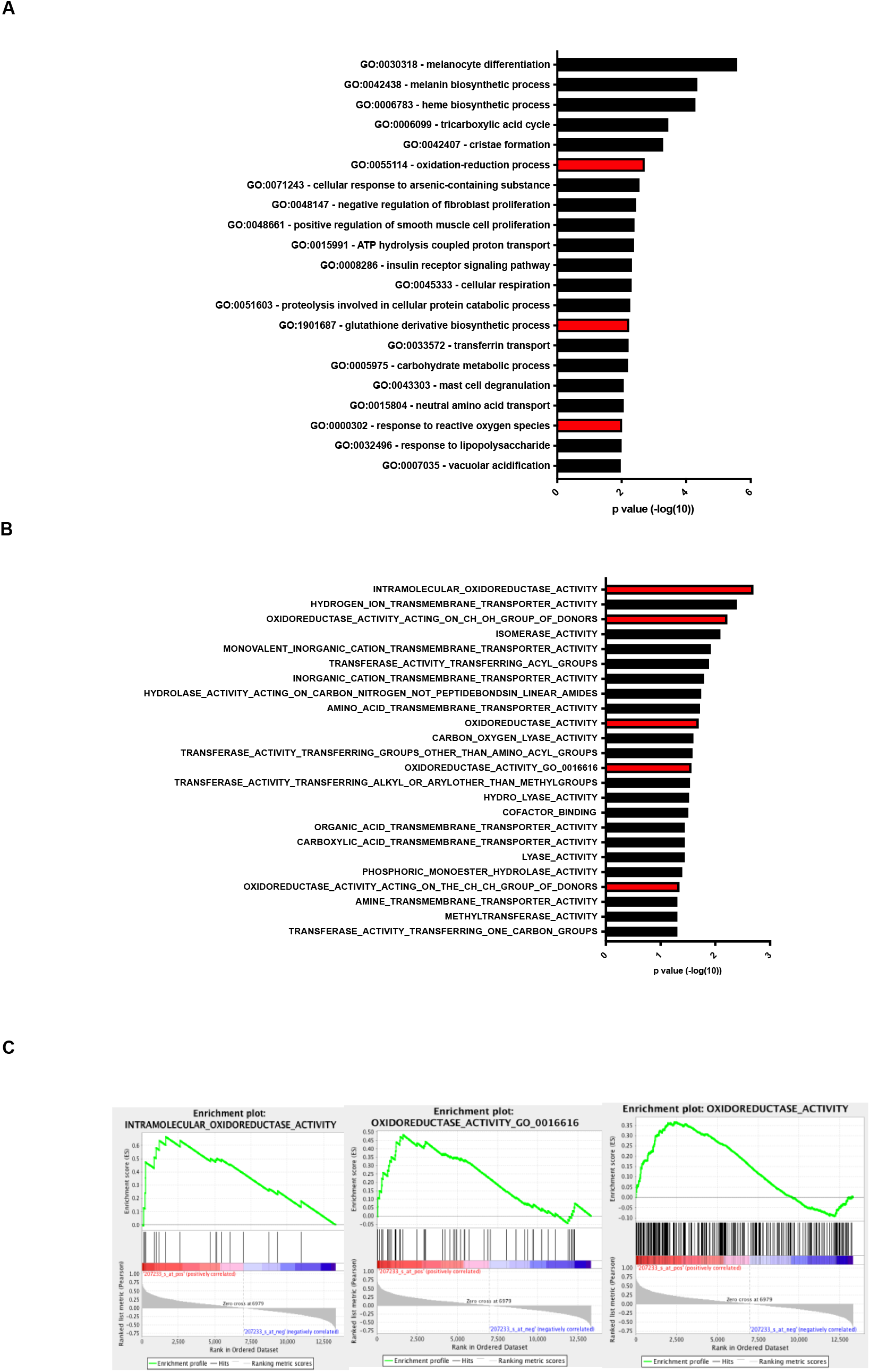
MITF correlates with oxidoreductase activities in multiple datasets. (**A**) Gene functional classification using DAVID revealed that multiple gene sets involved in redox regulation are significantly enriched among genes downregulated after MITF^†^ knockdown. (**B**) Gene set enrichment analysis using GO:Molecular function gene set database revealed that low MITF across 88 short-term melanoma cultures is significantly associated with decreased expression of genes with oxidoreductase activity. (**C**) Representative enrichment plots of oxidoreductase activity-related gene sets in high MITF melanomas compared to low MITF melanomas from 88 short-term cultured melanomas. † MITF: Microphthalmia-associated transcription factor

**Fig. S2.**
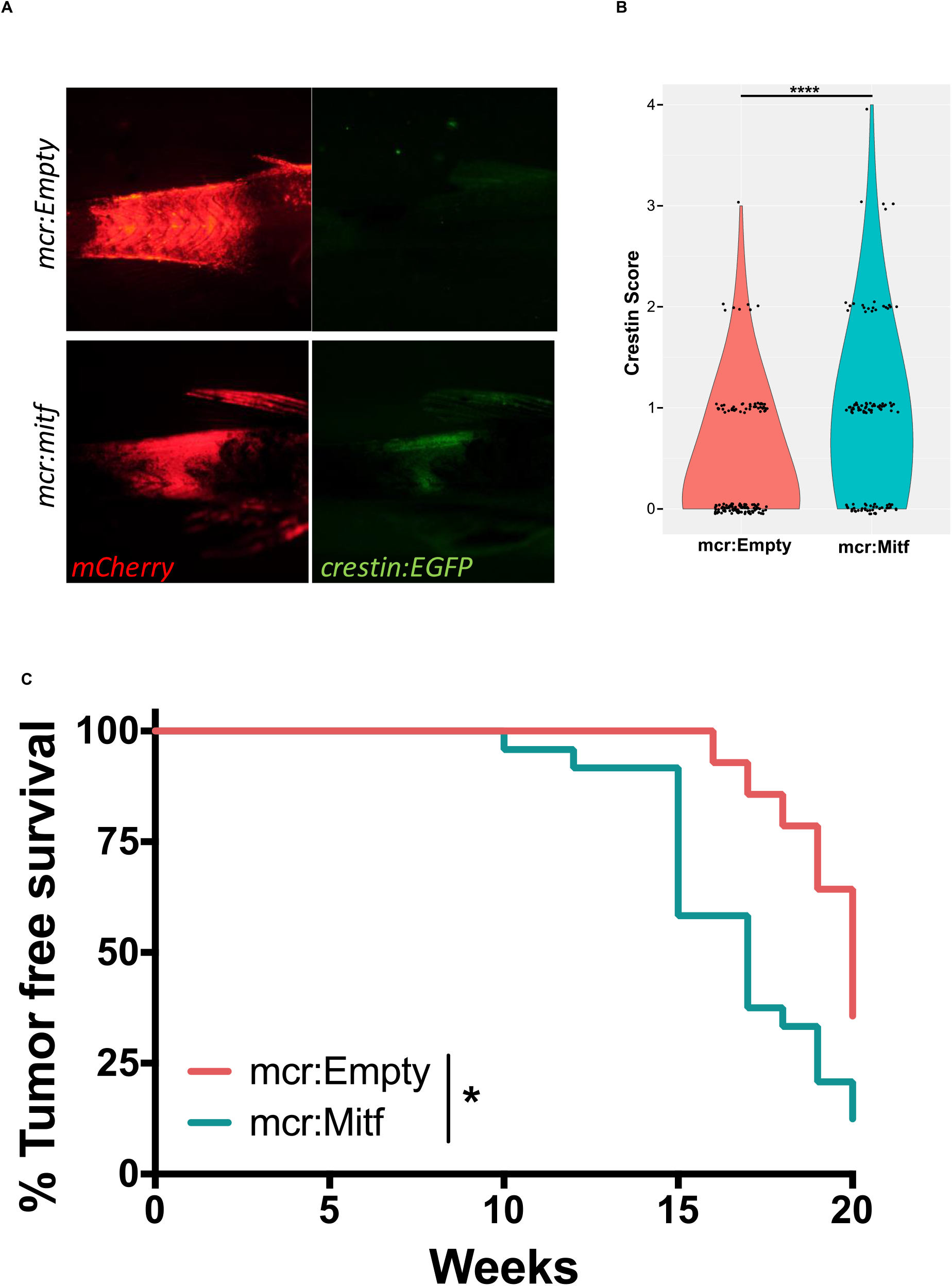
Features of zebrafish melanoma model. Fluorescent images (**A**) and quantitation (**B**) show rescued melanocytes (*mitf:mCherry*+ cells) and neural crest reactivation in early melanoma patches (*crestin:EGFP*+ cells) in 6-week-old *MITF*^†^ overexpressing (*mcr:MITF*) fish compared with control (*mcr:Empty*) fish (scored on a 0-5 scale, with 5 representing a tumor). (**C**) Kaplan-Meier curve depicting the percent of *mcr:MITF* fish with tumor-free survival compared to control *mcr:Empty* fish. † MITF: Microphthalmia-associated transcription factor

**Fig. S3.**
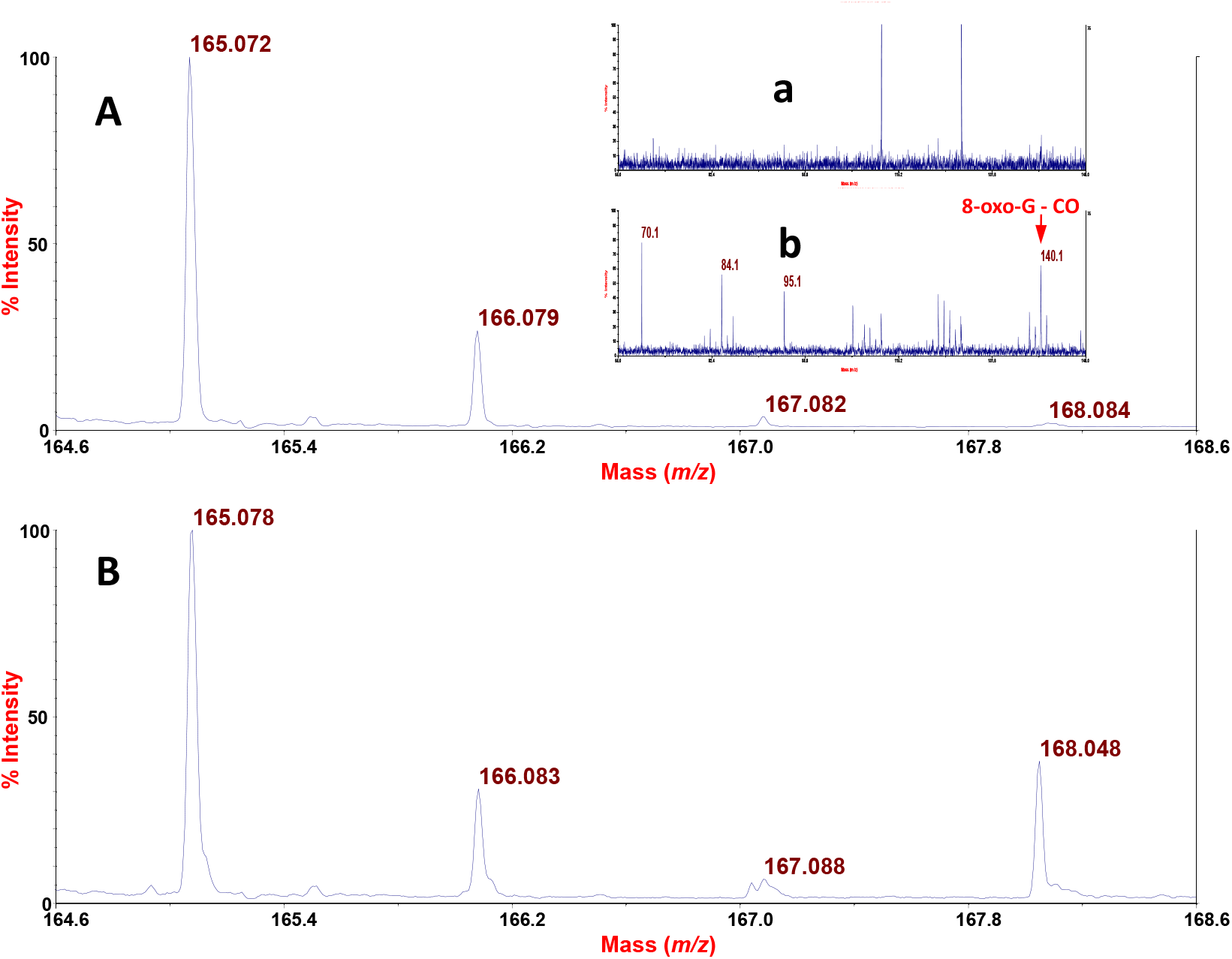
MALDI-TOF mass spectra. (**A**) CCA matrix; (**B**) DNA from Mitf1, showing a peak for 8-oxo-G at m/z 168.048; a corresponding TOF/TOF spectrum of m/z 168 gave a peak at m/z 140.1 from loss of CO (insets a and b correspond to A and B, respectively).

